# Neuromuscular denervation and deafferentation but not motor neuron death are disease features in the *Smn^2B/-^* mouse model of SMA

**DOI:** 10.1101/2022.04.21.489030

**Authors:** Maria J. Carlini, Marina K. Triplett, Livio Pellizzoni

## Abstract

Spinal muscular atrophy (SMA) is a neurodegenerative disease characterized by loss of motor neurons and skeletal muscle atrophy which is caused by ubiquitous deficiency in the survival motor neuron (SMN) protein. Several cellular defects contribute to sensory-motor circuit pathology in SMA mice, but the underlying mechanisms have often been studied in one mouse model without validation in other available models. Here, we used *Smn^2B/-^* mice to investigate specific behavioral, morphological, and functional aspects of SMA pathology that we previously characterized in the *SMNΔ7* model. *Smn^2B/-^* SMA mice on a pure FVB/N background display deficits in body weight gain and muscle strength with onset in the second postnatal week and median survival of 19 days. Morphological analysis revealed severe loss of proprioceptive synapses on the soma of motor neurons and prominent denervation of neuromuscular junctions (NMJs) in axial but not distal muscles. In contrast, no evidence of cell death emerged from analysis of several distinct pools of lumbar motor neurons known to be lost in the disease. Moreover, SMA motor neurons from *Smn^2B/-^* mice showed robust nuclear accumulation of p53 but lack of phosphorylation of serine 18 at its amino-terminal, which selectively marks degenerating motor neurons in the *SMNΔ7* mouse model. These results indicate that NMJ denervation and deafferentation, but not motor neuron death, are conserved features of SMA pathology in *Smn^2B/-^* mice.

## Introduction

Spinal muscular atrophy (SMA) is an inherited neurodegenerative disease characterized by loss of motor neurons and skeletal muscle atrophy, leading to motor dysfunction, paralysis and eventually death in its most severe form [1,2]. SMA is caused by ubiquitous reduction in the levels of the survival motor neuron (SMN) protein—reflecting homozygous loss of the *SMN1* gene but preservation of the nearly identical *SMN2* gene [3]. Due to inefficient splicing of exon 7 [4], the *SMN2* gene mainly produces an unstable protein isoform (SMNΔ7) and low levels of full-length functional SMN that cannot compensate for the loss of *SMN1*, leading to the disease [1,2].

Since higher *SMN2* copy numbers are associated with reduced disease severity in SMA patients, most therapeutic efforts have focused on increasing expression of SMN through modulation of *SMN2* splicing or SMN replacement by gene therapy [5–8]. Importantly, three distinct SMN-inducing therapies have demonstrated efficacy in clinical trials and are currently approved for treatment of SMA patients [9–14]. Nevertheless, these therapies are not a complete cure for the disease, and the development of additional therapeutics that could help address unmet clinical needs of SMA patients is necessary [15,16]. In principle, these novel drugs should target pathogenic events and enhance clinical benefit in combination with SMN-inducing therapies.

Loss of motor neurons is a hallmark of SMA pathology that is widely recognized to have a significant clinical impact on the disease course [1,2]. Genetic studies in mouse models have indicated that neurodegeneration in SMA is primarily an intrinsic, cell-autonomous process induced by SMN deficiency in motor neurons [17,18]. Moreover, motor neurons are the only cell type known to die during disease course, and their loss represents an irreversible pathogenic event that cannot be corrected after it has occurred. Thus, preventing motor neuron degeneration has important clinical implications for SMA therapy. The availability of distinct mouse models of SMA has been instrumental to the study of disease mechanisms and preclinical evaluation of SMA therapies that are now approved for treatment of patients [19–22]. To date, several non-mutually exclusive mechanisms have been proposed to contribute to motor neuron loss in SMA mice [23–29]. Our previous work has highlighted that motor neurons degenerate through activation of the tumor suppressor p53 in the *SMNΔ7* mouse model of SMA [27]. Importantly, not all motor neurons are equally susceptible to SMN deficiency [30], and we identified at least two distinct pathogenic events that converge on p53 to trigger selective death of vulnerable SMA motor neuron pools [27]: i) upregulation of p53; and ii) phosphorylation of the amino-terminal transcriptional activation domain of p53 including serine 18 (p53^S18^). Mechanistically, we showed that disruption of SMN-dependent alternative splicing of specific exons in the pre-mRNAs of Mdm2 and Mdm4 – two well-established inhibitors of p53’s stability and function – is responsible for nuclear accumulation of p53 in SMA motor neurons [28]. We also showed that U12 splicing-dependent dysregulation of the *Stasimon/Tmem41b* gene induced by SMN deficiency contributes to the cascade of events leading to p53 phosphorylation and death of motor neurons in *SMNΔ7* SMA mice [29,31]. Lastly, we found that Stasimon dysfunction induces p38 mitogen-activated protein kinase (p38MAPK) activation and that pharmacological inhibition of p38αMAPK reduces p53 phosphorylation and improves motor neuron survival in *SMNΔ7* mice [29], highlighting the neuroprotective effects of p38αMAPK inhibition in SMA mice. However, these as well as other proposed death mechanisms of SMA motor neurons have often been studied only in one mouse model without cross validation in other available models.

This study was designed as part of our efforts to determine whether p53 activation is a shared pathogenic mechanism associated with motor neuron death across different SMA models and to further validate pharmacological inhibition of p38αMAPK as a candidate neuroprotective approach. We used the *Smn^2B/-^* mouse model of SMA that harbors a hypomorphic *Smn* allele (*Smn^2B^*) with a mutation in the splicing regulatory sequence of exon 7 of the endogenous gene and a knockout *Smn* allele in a pure FVB/N genetic background [32]. We performed behavioral and morphological studies of sensory-motor circuit pathology in this mouse model to determine the effects of SMN deficiency on synaptic integrity and motor neuron survival. Consistent with previous studies [32], we found that *Smn^2B/-^* SMA mice display reduced weight gain, impaired motor function, and median survival of ~19 days. We also found severe loss of proprioceptive synapses on motor neurons and selective neuromuscular junction (NMJ) denervation of axial but not distal muscles. Surprisingly, however, we found no evidence for motor neuron loss, which correlated with nuclear accumulation of p53 but lack of amino-terminal phosphorylation of p53^S18^ in SMA motor neurons. Collectively, these findings highlight shared and distinct features of SMA pathology in *SMNΔ7* and *Smn^2B/-^* mice, which have important implications for guiding the selection of appropriate models for basic and translational studies of specific aspects of SMA pathology in the future. They also indicate that the *Smn^2B/-^* model is not well suited for *in vivo* testing of neuroprotective drugs that specifically target the motor neuron death pathway.

## Materials and methods

### Mouse lines

All mouse work was performed in accordance with the National Institutes of Health Guidelines on the Care and Use of Animals, complied with all ethical regulations and was approved by the IACUC committee of Columbia University. Mice were housed in a 12h/12h light/dark cycle with access to food and water *ad libitum*. Heterozygous *Smn^+/−^* mice harboring the *Smn1^tm1Msd^* knockout allele [33] on a pure FVB/N genetic background were obtained by crossing the *SMNΔ7* mouse line FVB.Cg-Grm7^Tg(SMN2)89Ahmb^ Smn1^tm1Msd^ Tg(SMN2*delta7)4299Ahmb/J (Jax stock #005025) with FVB/N mice until removal of the *SMNΔ7* and SMN2 transgenes. The *Smn*^2B/2B^ mice on a pure FVB/N genetic background were previously described [32]. *Smn^2B/2B^* mice were crossed with *Smn^+/−^* mice to generate *Smn^2B/-^* SMA mice and *Smn^2B/+^* littermates that were used as normal controls. Equal proportions of mice of both sexes were used and aggregated data are presented because gender-specific differences were not found.

### Genotyping

Genotyping was performed from tail DNA using a common forward primer (5’-GATGATTCTGACATTTGGGATG-3’) and specific reverse primers (5’-TGGCTTATCTGGAGTTTCACAA-3’) and (5’-GAGTAACAACCCGTCGGATTC-3’) for wild type *Smn* and *Smn1^tm1Msd^* knockout alleles, respectively [34]. The *Smn^2B^* allele was genotyped using forward (5’-AACTCCGGGTCCTCCTTCCT-3’) and reverse (5’-TTTGGCAGACTTTAGCAGGGC-3’) primers as previously described [32].

### Behavioral assays

Mice from all experimental groups were monitored daily for survival and weight from birth to weaning at 21 days. Righting reflex was assessed by placing the mouse on its back and measuring the time it took to turn upright on its four paws (righting time). Cut-off test time was 60 seconds. For each testing session, the test was repeated three times and the mean of the recorded times was calculated. For the hindlimb suspension test [32], the mouse was suspended by its hindlimbs from the rim of a cylindrical container with cushioning at the bottom. The time it took for a mouse to fall from the rim into the container was recorded with a cut-off time of 60 seconds. Mice able to climb back to the rim of the container during the test were scored as meeting the cut-off time. The test was repeated twice for each testing session and the mean of the recorded times was calculated.

### Antibodies and fluorescent probes

For western blot analysis, we used an anti-SMN mouse monoclonal antibody (BD Transd Lab, clone 8, #610646; 1:10,000), an anti-Tubulin mouse monoclonal antibody (Sigma, clone DM1A, #T9026; 1:50,000), and a HRP conjugated goat anti-mouse secondary antibody (Jackson #115-035-044; 1:10,000). For spinal cord immunohistochemistry, we used goat anti-ChAT (Millipore #AB144P; 1:100), guinea pig anti-VGluT1 (Covance, custom made; 1:5,000) [18], rabbit anti-p53 (Leica Novocastra #NCL-p53-CM5p; 1:1,000) and rabbit anti-phosporylated-p53^S15^ (Cell Signaling #9284, Lot: #15; 1:250) as primary antibodies. For muscle immunohistochemistry, we used guinea pig anti-Synaptophysin 1 (Synaptic Systems #101-004; 1:500), rabbit anti-Neurofilament M (Millipore #AB1987; 1:500), and Alexa Fluor™ 555 conjugated α-bungarotoxin (Invitrogen, #B35451; 1:500). Species-specific secondary antibodies coupled to Cy3 or Cy5 were used as appropriate (Jackson ImmunoResearch Laboratories, Inc; 1:250).

### Protein analysis

For Western blot analysis, mice were sacrificed and spinal cord collection was performed in a dissection chamber under continuous oxygenation (95%O_2_/5%CO_2_) in the presence of cold (~12°C) artificial cerebrospinal fluid (aCSF) containing 128.35mM NaCl, 4mM KCl, 0.58mM NaH_2_PO_4_, 21mM NaHCO_3_, 30mM D-Glucose, 1.5mM CaCl_2_, and 1mM MgSO_4_. Total protein extracts were generated by homogenization of spinal cords in SDS sample buffer (2% SDS, 10% glycerol, 5% β-mercaptoethanol, 60mM Tris-HCl pH 6.8, and bromophenol blue), followed by brief sonication and boiling. Proteins were quantified using the *RC DC*™ Protein Assay (Bio-Rad) and analyzed by SDS/PAGE on 12% polyacrylamide gels followed by Western blotting as previously described [35].

### Immunohistochemistry

Animals were anesthetized and transcardially perfused with 4% paraformaldehyde (PFA). The spinal cord and skeletal muscles were dissected and post-fixed in 4% PFA for 4 hours. For immunohistochemistry, the spinal cords were briefly washed with PBS, specific lumbar segments were identified by the ventral roots and subsequently embedded in warm 5% agar. Transverse sections (75μm) of the entire spinal segment were obtained with a VT1000 S vibratome (Leica). All the sections were then blocked with 10% normal donkey serum in 0.01M PBS containing 0.4% Triton X-100 (PBS-T; pH 7.4) for 1 hour and incubated overnight at room temperature with different combinations of primary antibodies diluted in PBS-T. The following day, six washing steps of 10 minutes each were done prior to incubation with secondary antibodies for 3 hours in PBS. Another six washing steps were performed before sections were mounted in 30% glycerol/PBS. For NMJ analysis, skeletal muscles were cryoprotected through sequential immersion in 10% and 20% sucrose/0.1M phosphate buffer (PB) for 1 hour at 4°C followed by overnight immersion in 30% sucrose/0.1M PB at 4°C. The following day muscles were frozen embedded in Optimal Cutting Temperature (OCT) compound (Fisher), frozen on dry ice, and stored at −80°C until processing. Longitudinal cryosections (30μm) were collected onto Superfrost Plus glass slides (Fisher) using a CM3050S cryostat (Leica). Sections were washed once with PBS for 5 minutes to remove OCT, blocked for 1 hour with 5% donkey serum in TBS containing 0.2% Triton-X at room temperature and incubated with primary antibodies in blocking buffer overnight at 4°C. Following incubation, sections were washed three times for 10 minutes in TBS containing 0.2% Triton-X and then incubated with tetramethylrhodamine-conjugated α-bungarotoxin (Invitrogen #T1175, 1:500) and the appropriate secondary antibodies for 1 hour at room temperature, followed by 3 washing steps. Slides were mounted using Fluoromount-G Mounting Medium (SouthernBiotech).

### Confocal microscopy and image analysis

All images were collected with an SP5 confocal microscope (Leica) running the LAS AF software (v2.5.2.6939) and analyzed off-line using the Leica LAS X software (v1.9.0.13747). For motor neuron number quantification, 1024 × 1024 pixels images were acquired from all the 75μm sections of each specific spinal segment using a 20X objective at 3μm steps in the z-axis and a 200 Hz acquisition rate. Only motor neurons (ChAT^+^) with a clearly identifiable nucleus were counted to avoid double counting from adjoining sections. For quantification of VGluT1^+^ synapses, 1024 × 1024 pixels images were acquired from L2 spinal sections (75μm) using a 40X objective at 0.3μm steps in the z-axis and a 200 Hz acquisition rate. The total number of VGluT1^+^ synapses on soma was determined by counting all the corresponding inputs on the surface of each ChAT^+^ motor neuron cell body. At least 10 motor neurons per mouse were quantified. For NMJ analysis, 1024 × 1024 pixels images were acquired from 30μm muscle sections using a 20X objective at 2μm steps in the z-axis and a 200 Hz acquisition rate. Maximum intensity projections of confocal stacks were analyzed and at least 200 randomly selected NMJs per muscle were quantified. NMJs lacking any coverage of the α-bungarotoxin-labeled postsynaptic endplate by the presynaptic markers Synaptophysin and Neurofilament-M were scored as denervated.

### Statistical analysis

Statistical analysis was performed by two-tailed unpaired Student’s *t*-test or by two-way ANOVA followed by the Bonferroni’s multiple comparison test as indicated. Comparison of survival curves was performed using the Log-rank (Mantel-Cox) test. GraphPad Prism (v9.3.1) was used for all statistical analyses and P values are indicated as follows: *P<0.05; **P<0.01; ***P<0.001; **** P< 0.0001.

## Results

### Behavioral characterization of *Smn^2B/-^* SMA mice

*Smn^2B^* is a hypomorphic allele harboring a mutation in the splicing regulatory sequence of exon 7 in the mouse *Smn* gene leading to exon skipping and reduced expression of full-length Smn protein [22]. Accordingly, we first analyzed the levels of Smn expression from the *Smn^2B^* allele by Western blot analysis of spinal cord tissue and found that homozygous *Smn^2B/2B^* mice express half the levels of Smn protein relative to wild type (*Smn^+/+^*) mice at P16 (Fig S1A). A similar reduction in the levels of Smn expression was also found by comparing spinal cords from *Smn^2B/-^* SMA mice and control *Smn^2B/+^* littermates at P16 (Fig S1B). Thus, the *Smn^2B^* allele expresses approximately 25% of the amount of Smn protein relative to the wild type *Smn* allele in the mouse spinal cord, which is consistent with previous studies [22].

Next, we sought to characterize the disease phenotype of *Smn^2B/-^* SMA mice. We monitored daily weight gain from birth to weaning in *Smn^2B/-^* SMA mice and *Smn^2B/+^* littermates, which were used as normal controls in this study. While both groups displayed a similar weight gain in the first two postnatal weeks, *Smn^2B/-^* SMA mice showed a significant and progressive decline in weight relative to *Smn^2B/+^* littermates starting at P15 (Fig 1A). The decline in body weight was rapidly followed by death of *Smn^2B/-^* SMA mice, which displayed a median lifespan of 19 days (P < 0.0001, Log-rank Mantel-Cox test) (Fig 1B). To analyze motor function, we performed the righting reflex and the hindlimb suspension tests, which are two behavioral assays widely used to monitor disease-related motor phenotypes in mouse models of SMA. *Smn^2B/-^* SMA mice showed a comparable ability to acquire the righting reflex relative to controls during early postnatal development (Fig 1C). On the other hand, while *Smn^2B/+^* control mice rapidly improved and then maintained their performance in the hindlimb suspension test from P11 onward, *Smn^2B/-^* SMA mice not only failed to show improvement but also progressively worsened their performance over time, pointing to compromised hindlimb muscle strength (Fig 1D). Thus, consistent with previous studies [22,32], we found that *Smn^2B/-^* SMA mice display impaired weight gain and motor function as well as reduced survival.

**Fig 1.**
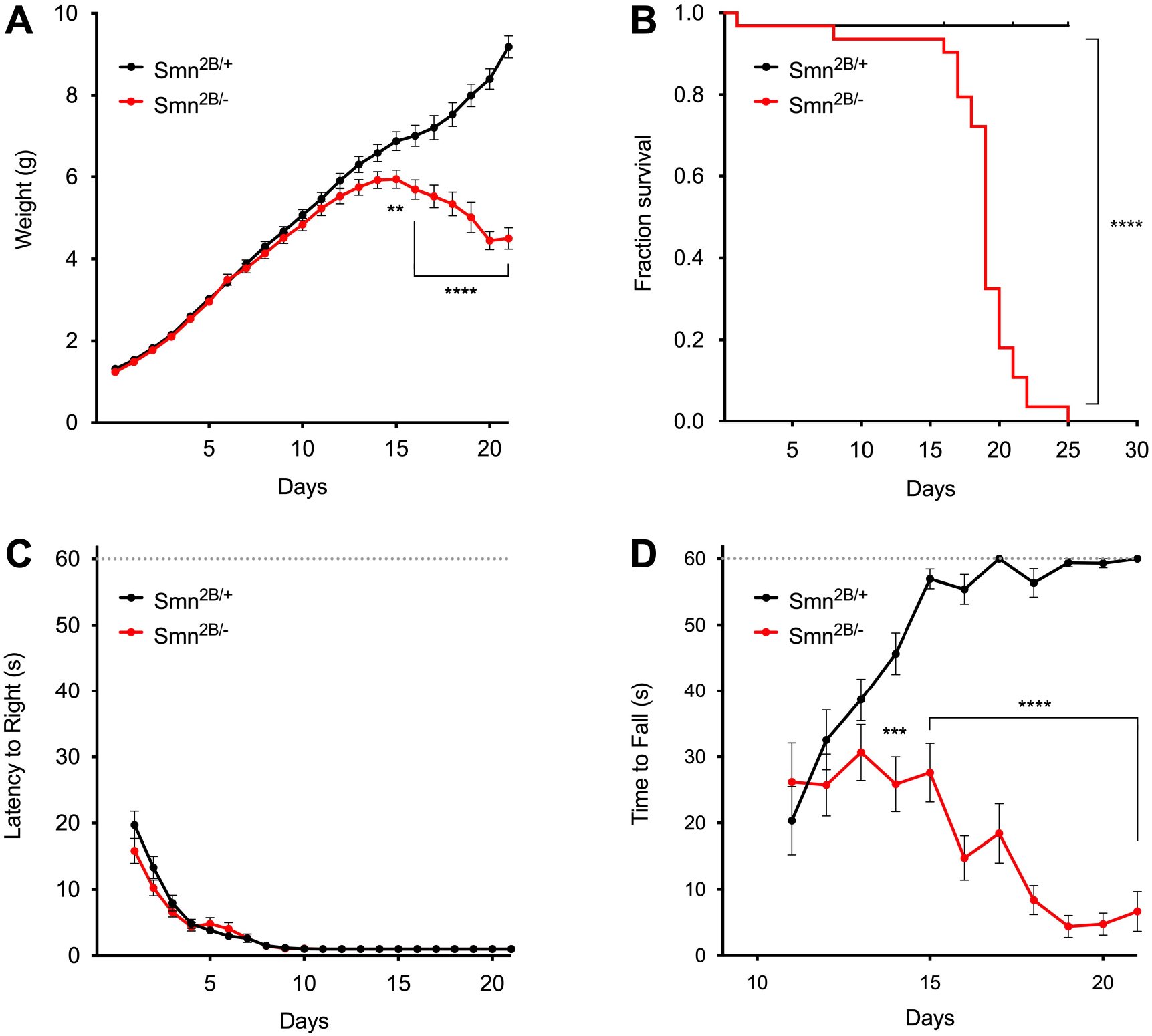
Behavioral characterization of *Smn^2B/-^* SMA mice. (A) Body weight of control *Smn^2B/+^* (n=32) mice and *Smn^2B/-^* SMA mice (n=31). Data represent mean and SEM. Statistics were performed with twoway ANOVA and Bonferroni’s multiple comparison test. ** P < 0.01; **** P < 0.0001. (B) Kaplan-Meier survival curves from the same experimental groups as in (A). Statistics were performed with Log-rank (Mantel-Cox) test. **** P < 0.0001. (C) Righting time from the same experimental groups shown in (A). Data represent mean and SEM. Statistics were performed with two-way ANOVA and Bonferroni’s multiple comparison test. Not Significant. (D) Time to fall in the hindlimb suspension test from the same experimental groups shown in (A). Data represent mean and SEM. Statistics were performed with two-way ANOVA and Bonferroni’s multiple comparison test. *** P < 0.001; **** P < 0.0001.

### Severe loss of proprioceptive synapses on motor neurons of *Smn^2B/-^* SMA mice

Spinal sensory-motor circuit dysfunction is one of the earliest pathological features of SMA in mouse models [1]. In particular, the loss of VGluT1^+^ proprioceptive synapses on the soma and proximal dendrites of motor neurons has been shown to occur early in the disease course [18,30,36,37] as well as independently from motor neuron loss in the *SMNΔ7* mouse model of SMA [18,27,28]. Furthermore, the molecular defects induced by Smn deficiency that contribute to deafferentation of SMA motor neurons have recently emerged, including dysregulation of U12 splicing of the *Stasimon* gene [29,38]. Since most of these studies were performed in the *SMNΔ7* mouse model of SMA, we sought to investigate the effects of SMN deficiency on the connectivity of proprioceptive synapses onto motor neurons in *Smn^2B/-^* SMA mice. As in our previous studies, we focused on lumbar motor neurons innervating disease-relevant proximal muscles and employed immunohistochemistry and confocal microscopy to quantify the number of VGluT1^+^ proprioceptive synapses juxtaposed to the soma of ChAT^+^ motor neurons. (Fig 2A). We found that the number of proprioceptive synapses onto motor neurons of *Smn^2B/-^* SMA mice is markedly reduced relative to *Smn^2B/+^* mice at P16 (Fig 2B). Thus, Smn deficiency induces severe deafferentation of motor neurons in the *Smn^2B/-^* mouse model of SMA.

**Fig 2.**
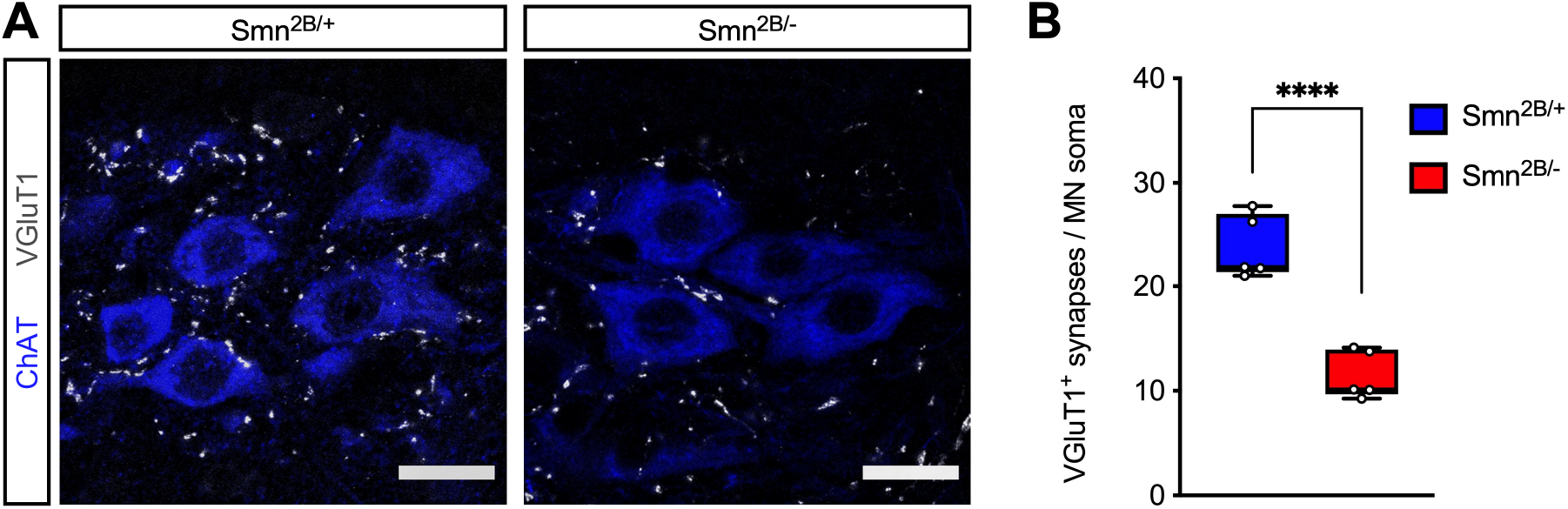
*Smn^2B/-^* SMA mice display severe loss of proprioceptive synapses on motor neurons. (A) Immunostaining of VGluT1^+^ synapses (grayscale) and ChAT^+^ motor neurons (blue) in the L2 spinal cord from control (*Smn^2B/+^*) and SMA (*Smn^2B/-^*) mice at P16. Scale bars: 25μm. (B) Number of VGluT1^+^ synapses on the soma of L2 motor neurons from the same groups as in (A) at P16. The box-and-whiskers graph shows the individual values, median, interquartile range, minimum and maximum from 5 mice per group. Statistics were performed with two-tailed unpaired Student’s t-test. **** P < 0.0001.

### Loss of neuromuscular junction innervation in axial but not distal muscles of *Smn^2B/-^* SMA mice

The loss of neuromuscular junction (NMJ) synapses between motor neurons and skeletal muscle is a key disease feature of SMA pathology with proximal and axial muscles being more affected than distal muscles in both patients and mouse models [39–42]. Therefore, we sought to examine NMJ innervation in the axial muscle *quadratus lumborum* (QL) and the distal muscle *tibialis anterior* (TA) in *Smn^2B/-^* SMA mice relative to *Smn^2B/+^* controls at P16. To do so, we performed NMJ staining using antibodies against Neurofilament-M and Synaptophysin as pre-synaptic markers and α-bungarotoxin as post-synaptic marker of the motor endplate. As expected, 100% of the NMJs were fully innervated in both QL and TA muscles of control mice (Fig 3). Importantly, analysis of the QL muscle from *Smn^2B/-^* SMA mice revealed strong NMJ denervation as many α-bungarotoxin-labeled motor endplates completely lacked pre-synaptic coverage by nerve terminals (Fig 3A). Quantification of this defect showed that approximately 40% of the NMJs in QL muscle are denervated in *Smn^2B/-^* SMA mice (Fig 3B). In contrast, nearly all the NMJs in the TA muscle from *Smn^2B/-^* SMA mice were innervated (Fig 3D). Taken together, these results demonstrate marked and preferential loss of NMJ innervation from an axial muscle relative to a distal muscle in *Smn^2B/-^* SMA mice, which is consistent with the features of neuromuscular pathology in the human disease.

**Fig 3.**
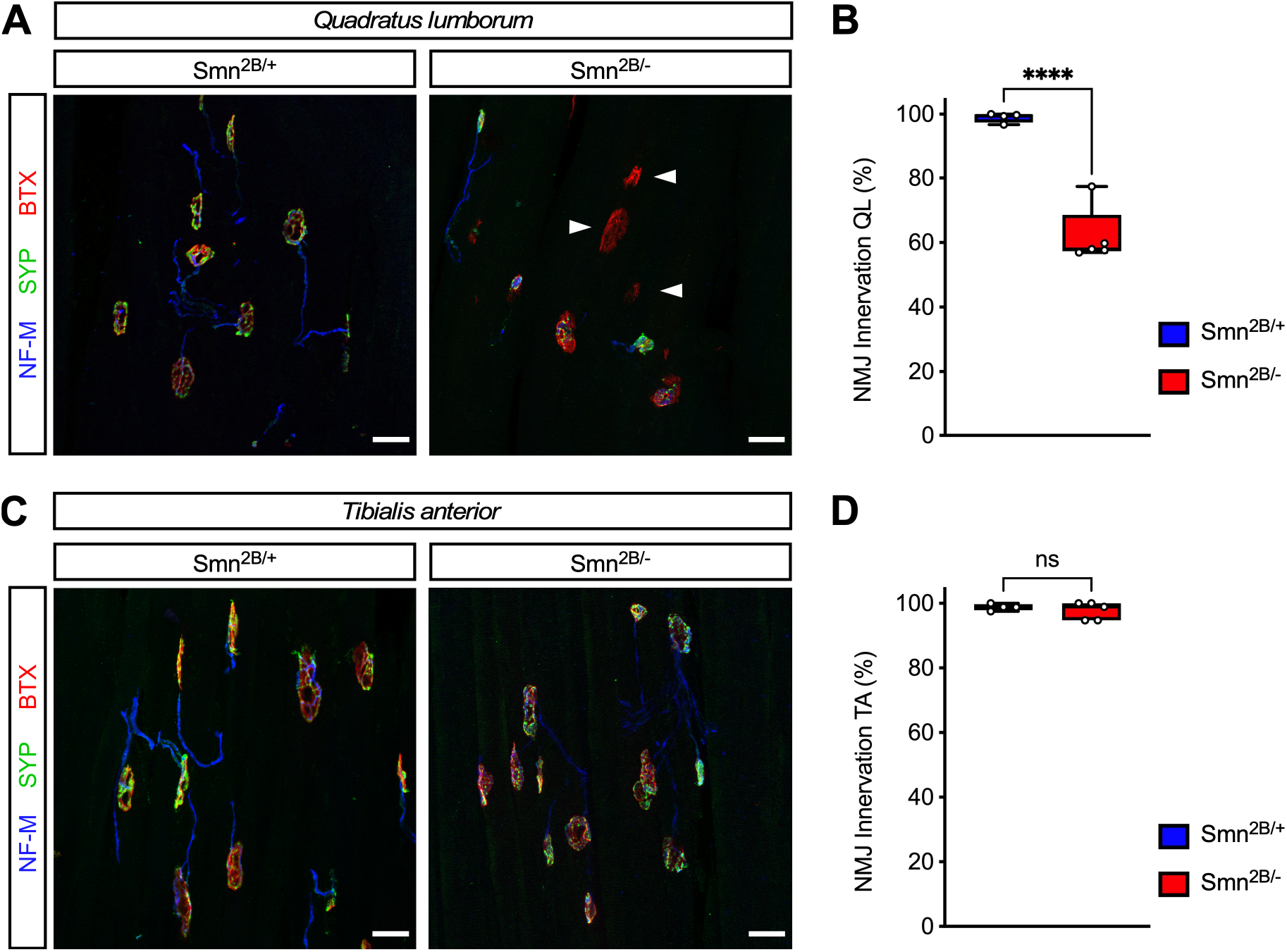
Selective loss of NMJ innervation in axial but not distal muscles in *Smn^2B/-^* SMA mice. (A and C) NMJ staining with bungarotoxin (BTX), Synaptophysin (SYP), and Neurofilament-M (NF-M) of the axial muscle *quadratus lumborum* (A) and the distal muscle *tibialis anterior* (C) from control (*Smn^2B/+^*) and SMA (*Smn^2B/-^*) mice at P16. Arrowheads indicate denervated NMJs. Scale bars: 25 μm. (B and D) Percentage of fully innervated NMJs from the same groups as in (A) and (C). The box-and-whiskers graph shows the individual values, median, interquartile range, minimum and maximum from *Smn^2B/+^* (n=4) and *Smn^2B/-^* (n=5) mice. Statistics were performed with two-tailed unpaired Student’s t-test. **** P < 0.0001, ns=not significant.

### Survival of motor neurons is not affected in *Smn^2B/-^* SMA mice

The selective degeneration of specific pools of motor neurons is a hallmark of SMA [1,2]. Reflecting the characteristic profile of differential muscle vulnerability, motor neuron pools innervating proximal and axial muscles are more prominently affected and preferentially lost in the disease. Accordingly, vulnerable pools comprise lumbar motor neurons residing in the L1 and L2 segments of the spinal cord as well as in the L5 medial motor column (MMC), which innervate proximal and axial muscles [18,30,41]. In contrast, L5 lateral motor column (LMC) motor neurons that innervate distal muscles are resistant to death in SMA. We sought to determine whether motor neurons in the *Smn^2B/-^* SMA mice displayed a similar profile of differential vulnerability to death induced by SMN deficiency. To do so, we performed immunostaining of all sections from the L1, L2 and L5 segments of the spinal cord with antibodies against ChAT and counted the total number of ChAT^+^ motor neurons in each of these segments from *Smn^2B/-^* SMA mice and *Smn^2B/+^* control littermates at P16 (Fig. 4A, 4C and 4E). Surprisingly, we found no significant loss of motor neurons in any of the lumbar segments analyzed from the spinal cord of *Smn^2B/-^* mice relative to controls (Fig. 4B, 4D, 4F and 4G). This analysis indicates that SMN deficiency does not affect the survival of motor neurons in the *Smn^2B/-^* mouse model of SMA at a late symptomatic time point.

**Fig 4.**
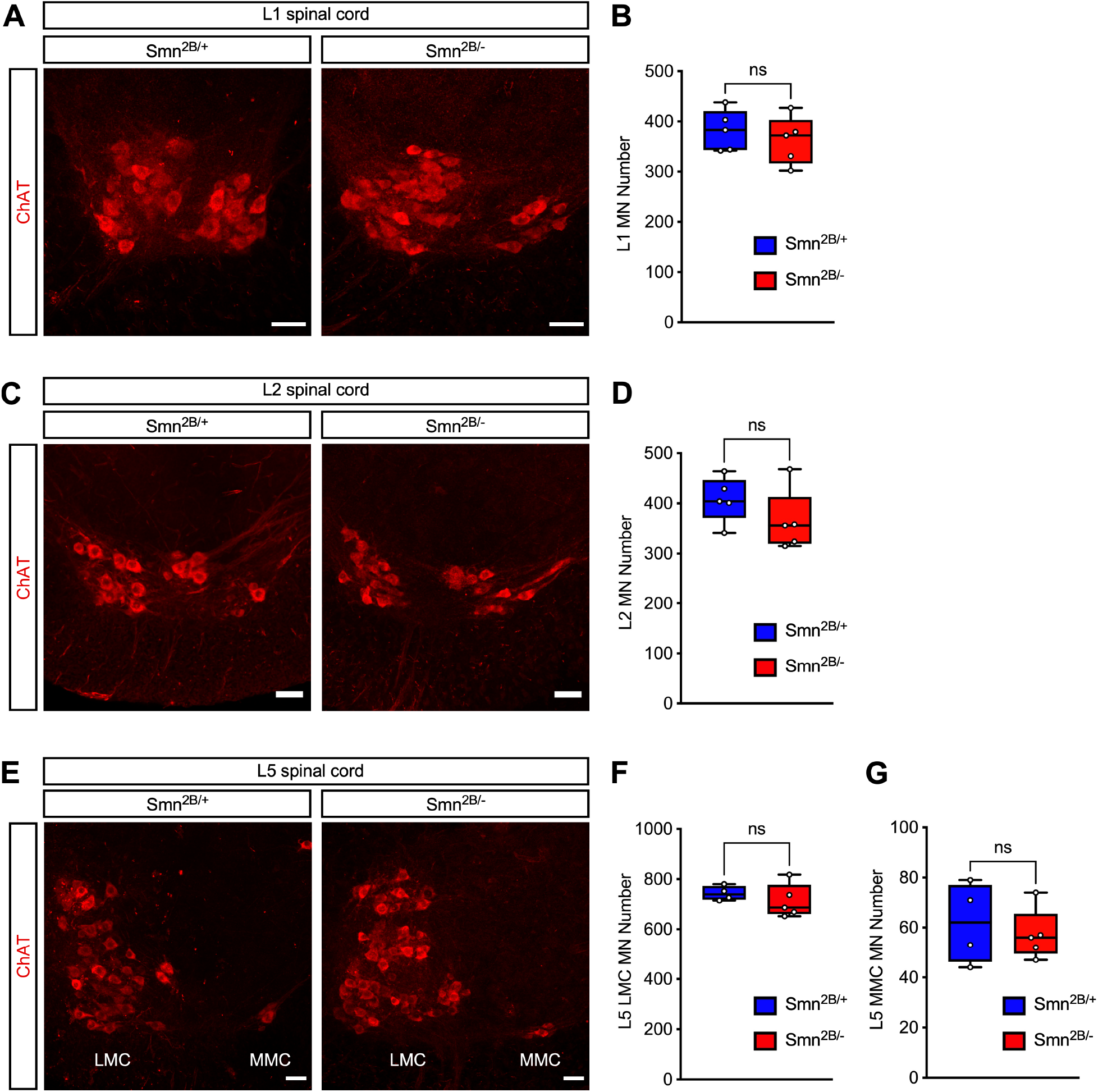
Death of lumbar motor neurons is not a disease feature of *Smn^2B/-^* SMA mice. (A) ChAT immunostaining of motor neurons in the L1 spinal segment from control (*Smn^2B/+^*) and SMA (*Smn^2B/-^*) mice at P16. Scale bars: 50μm. (B) Total number of L1 motor neurons from the same groups as in (A). The box- and-whiskers graph shows the individual values, median, interquartile range, minimum and maximum from 5 mice per group. Statistics were performed with two-tailed unpaired Student’s t-test. ns=not significant. (C) ChAT immunostaining of motor neurons in the L2 spinal segment from control (*Smn^2B/+^*) and SMA (*Smn^2B/-^*) mice at P16. Scale bars: 50μm. (D) Total number of L2 motor neurons from the same groups as in (C). The box-and-whiskers graph shows the individual values, median, interquartile range, minimum and maximum from 5 mice per group. Statistics were performed with two-tailed unpaired Student’s t-test. ns=not significant. (E) ChAT immunostaining of motor neurons in the L5 spinal segment from control (*Smn^2B/+^*) and SMA (*Smn^2B/-^*) mice at P16. Scale bars: 50μm. (F and G) Total number of L5 LMC (F) and L5 MMC (G) motor neurons from the same groups as in (E). The box-and-whiskers graph shows the individual values, median, interquartile range, minimum and maximum from *Smn^2B/+^* (n=4) and *Smn^2B/-^* (n=5) mice. Statistics were performed with two-tailed unpaired Student’s t-test. ns=not significant.

### SMN deficiency induces p53 accumulation but not serine 18 phosphorylation in motor neurons of *Smn^2B/-^* SMA mice

Our previous studies implicated activation of a p53-dependent pathway in the selective death of motor neurons in *SMNΔ7* SMA mice [27–29]. Therefore, we sought to investigate this pathway in *Smn^2B/-^* SMA mice, which do not display significant loss of motor neurons. As for the analysis of motor neuron survival, we focused on the study of the L1, L2 and L5 segments of the spinal cord from *Smn^2B/-^* SMA mice and *Smn^2B/+^* controls at P16. First, we performed immunohistochemistry experiments with antibodies against total p53 as well as ChAT to identify motor neurons followed by confocal microscopy. These experiments revealed strong nuclear accumulation of p53 in L1 (Fig 5A and 5B), L2 (Fig S2A and S2B) and L5 LMC and MMC (Fig S3A and S3B) motor neurons from *Smn^2B/-^* SMA mice but not from control *Smn^2B/+^* mice. Quantification showed nuclear accumulation of p53 in approximately 40% of vulnerable L1, L2 and L5 MMC motor neurons as well as resistant L5 LMC motor neurons from *Smn^2B/-^* SMA mice (Fig 5D, Fig S2D, Fig S3E and S3G). Moreover, we found nuclear p53 immunoreactivity in other spinal cells from *Smn^2B/-^* mice (Fig 5A, Fig S2A and S3A). Thus, SMN deficiency induces robust p53 accumulation in both motor neurons and other spinal cells in the *Smn^2B/-^* mouse model of SMA.

**Fig 5.**
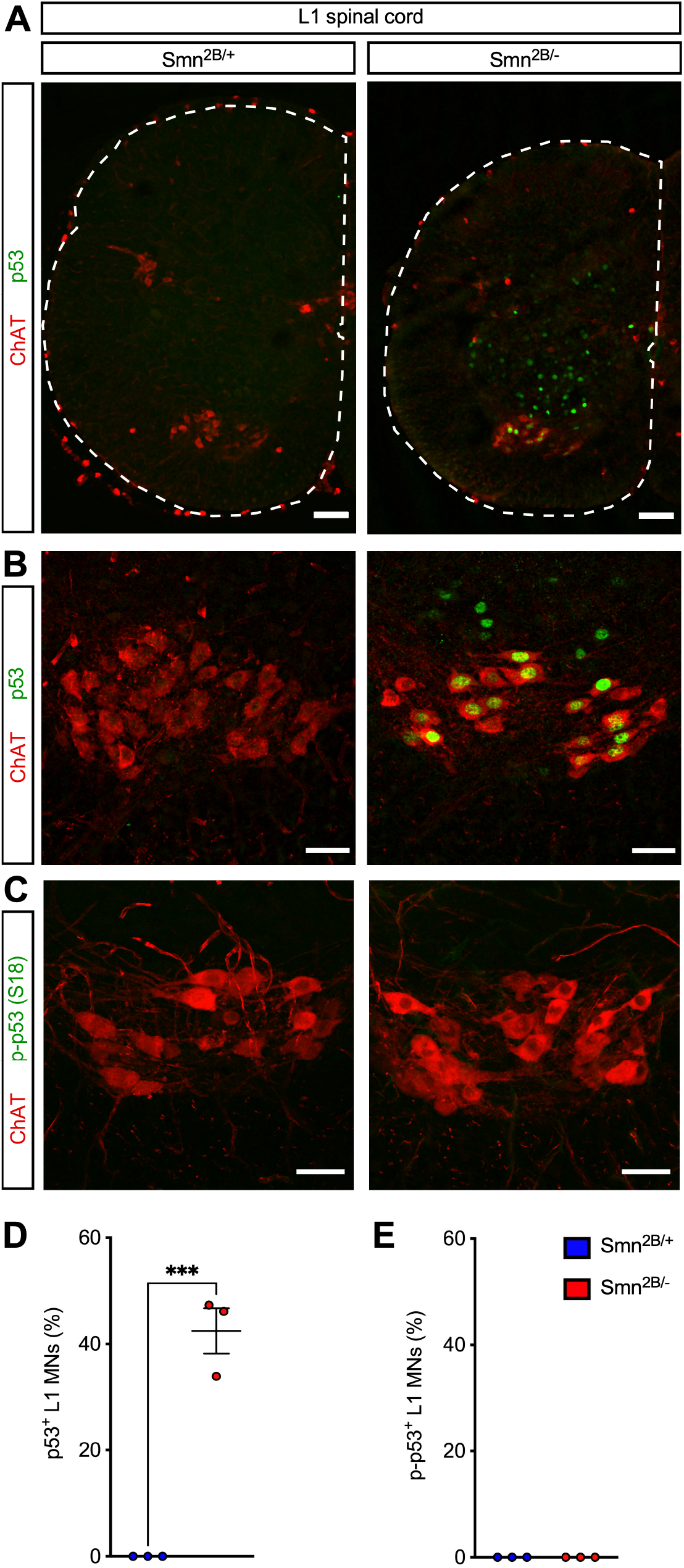
Smn deficiency induces p53 accumulation but not serine 18 phosphorylation in L1 motor neurons of *Smn^2B/-^* SMA mice. (A) ChAT and p53 immunostaining of the L1 spinal cord from control (*Smn^2B/+^*) and SMA (*Smn^2B/-^*) mice at P16. Scale bars: 100 μm. (B) ChAT and p53 immunostaining of L1 motor neurons from the same groups as in (A). Scale bars: 50 μm. (C) ChAT and phospho-p53^S18^ immunostaining of L1 motor neurons from the same groups as in (A). Scale bars: 50 μm. (D) Percentage of p53^+^ L1 motor neurons from the same groups as in (A). (E) Percentage of phospho-p53^S18+^ L1 motor neurons from the same groups as in (A). Data represents individual values, mean and SEM from 3 mice per group. Statistics were performed with two-tailed unpaired Student’s t-test. *** P < 0.001.

We previously showed that p53 nuclear accumulation is necessary but not sufficient to induce motor neuron death [27,28], which additionally requires phosphorylation of several serine residues in the amino terminus of p53 [27,29]. We also showed that phosphorylation of serine 18 of p53 (phospho-p53^S18^) selectively marks motor neurons destined to degenerate in SMA mice but is absent from resistant SMA neurons exhibiting p53 accumulation at late stages of disease [27,29]. Therefore, we investigated the expression of phospho-p53^S18^ by immunostaining of the L1, L2 and L5 spinal segments from *Smn^2B/-^* SMA mice and *Smn^2B/+^* controls at P16. Noteworthy, we did not detect any immunostaining of phospho-p53^S18^ in SMA motor neurons from *Smn^2B/-^* SMA mice (Fig 5C and 5E, Fig S2C and S2E, Fig S3C, S3D, S3F and S3H). Thus, despite strong nuclear accumulation of p53, the absence of detectable expression of phospho-p53^S18^ correlates with the lack of motor neuron loss in the *Smn^2B/-^* mouse model of SMA.

## Discussion

Here we carried out behavioral and morphological characterization of SMA pathology in the *Smn^2B/-^* mouse model of SMA. By monitoring the same parameters of sensory-motor circuit pathology and using the same assays we previously employed in our studies of *SMNΔ7* SMA mice, the study design allows direct comparison of our findings in the two models. Accordingly, we document similar features of synaptic pathology in *Smn^2B/-^* and *SMNΔ7* SMA mice, including severe loss of proprioceptive synapses on the soma of motor neurons and selective NMJ denervation of axial but not distal muscles. Surprisingly, however, we report the lack of significant loss of lumbar motor neurons at a late symptomatic stage of disease in *Smn^2B/-^* SMA mice that is in stark contrast with findings in *SMNΔ7* SMA mice. The observed differences in motor neuron survival are consistent with our proposed mechanisms of motor neuron death in *SMNΔ7* SMA mice implicating both nuclear accumulation and amino-terminal phosphorylation of p53 [27–29], the latter of which does not occur in *Smn^2B/-^* SMA mice. Collectively, these findings highlight shared and distinct features of SMA pathology across mouse models of SMA and indicate that *Smn^2B/-^* SMA mice are suitable for the study of some but not all the aspects of sensory-motor circuit pathology. Moreover, the lack of motor neuron death hinders the use of this model for *in vivo* testing of neuroprotective drugs specifically aimed at targeting motor neuron death.

The results of behavioral analysis of the SMA phenotype in *Smn^2B/-^* mice are well aligned with previous studies in the same model [32]. Accordingly, we found that *Smn^2B/-^* mice display a decline in weight gain at about two weeks of age that is mirrored by signs of progressive muscle weakness as revealed by failure to perform in the hindlimb suspension test. These deficits are compounded by shortened lifespan with a median survival of 19 days. Interestingly, however, *Smn^2B/-^* mice acquire the ability to right themselves in a manner indistinguishable from control littermates, which is very different from the severe impairment in performing this motor function found in *SMNΔ7* mice [17,30]. The reason for this difference in motor behavior remains to be established but could relate to a later onset in the loss of proprioceptive synapses in the *Smn^2B/-^* mice [37] at a time when the contribution of these synapses to the righting behavior is outweighed by the activity of descending vestibulo-spinal pathways [43,44].

We show here that the number of VGluT1^+^ excitatory synapses of proprioceptive neurons on the soma of motor neurons is severely reduced in *Smn^2B/-^* mice. Deafferentation of motor neurons was previously reported as one of earliest synaptic defects occurring in the *SMNΔ7* mouse model [30], mainly resulting from the effects of SMN deficiency in proprioceptive neurons [18]. Consistent with our results, loss of proprioceptive synapses on motor neurons was also observed in *Smn^2B/-^* mice on a different genetic background (C57BL/6) as well as in the *Taiwanese* model of SMA [37,45–47]. Thus, motor neuron deafferentation emerges as a conserved pathogenic event across all mouse models of SMA. Through rescue experiments in *SMNΔ7* mice, we have previously shown the direct contribution of U12 splicing dysregulation and Stasimon dysfunction in this process [29,31,38]. Other studies using *Taiwanese* SMA mice implicated deficits in pathways related to UBA1/GARS and Plastin [46,47]. Moreover, activation of the classical complement cascade has been linked to the execution of synaptic elimination of proprioceptive synapses in *SMNΔ7* mice [48]. It remains to be established whether these findings can be reconciled into a coherent cascade of events and the same mechanisms are responsible for the loss of central synapses in the different mouse models.

Our analysis of neuromuscular pathology reveals strong loss of NMJ innervation in the axial muscle QL but nearly complete sparing of the distal muscle TA. These findings complement and extend previous studies of NMJ pathology in *Smn^2B/-^* mice [22,37,49], indicating that the QL is among the most severely affected muscles in this mouse model. They are also consistent with the preferential susceptibility to NMJ denervation of proximal and axial SMA muscles that is observed across mouse models and appears more pronounced in *SMNΔ7* mice [37,40,50,51]. Lastly, the loss of NMJs from the QL muscle without death of the corresponding innervating motor neurons corroborates the conclusion that these two key pathogenic events are mechanistically uncoupled in SMA [28,29,52–54].

A surprising finding of this study is the lack of motor neuron loss, which is a hallmark of SMA pathology [1,2]. We investigated distinct pools of lumbar motor neurons known to be highly vulnerable in *SMNΔ7* mice [18,27,30,41], which include L1 and L2 as well as L5 MMC motor neurons. However, in all instances we found no significant reduction in the total number of spinal motor neurons at a late symptomatic time point (P16) in *Smn^2B/-^* mice, highlighting a marked difference between *SMNΔ7* and *Smn^2B/-^* mice. These results disagree with earlier studies reporting loss of motor neurons already at P11 in *Smn^2B/-^* mice on an FVB/N background that are identical to those analyzed here [32]. The reason for the discrepancy is unclear, but one possibility may lie in the accuracy of estimating motor neurons by sampling a subset of sections [32,55,56] as compared to counting the total number of motor neurons in all sections from the entire spinal segment (this study and [18,27–30,37]), the latter of which we consider more reliable. Other potentially confounding elements relate to the methods used for identification of specific spinal segments and the pooling of motor neuron counts obtained from sections spanning multiple segments that differ in the overall number of motor neurons as well as their susceptibility to disease [32,55,56]. Along these lines, we note that significant loss of motor neurons was also reported to occur at P15-P16 in *Smn^2B/-^* mice on the C57BL/6 background [32,55,56], but a recent study using the same experimental approach employed here found only limited loss of L1 motor neurons at P26, which is two days past median survival, but neither at earlier time points nor in other spinal segments [37]. Therefore, although we cannot exclude the possibility that a small loss of motor neurons may occur in *Smn^2B/-^* mice on the FVB/N background at times beyond their median survival, we conclude that motor neuron death is not a disease-relevant feature of SMA pathology in *Smn^2B/-^* mouse models.

To identify a potential reason for the lack of motor neuron loss in *Smn^2B/-^* mice, we investigated the status of p53 expression that we have previously linked to the death pathway in *SMNΔ7* mice [27]. By immunostaining experiments, we found widespread nuclear accumulation of p53 in SMA motor neurons from *Smn^2B/-^* mice at P16. Similar to the situation we reported in *SMNΔ7* mice at late symptomatic stages [27,28], p53 accumulation in *Smn^2B/-^* mice is observed in vulnerable L1, L2 and L5 MMC motor neurons as well as resistant L5 LMC motor neurons and other spinal cells that do not degenerate in the disease. These findings are consistent with the transcriptional upregulation of some p53-regulated genes in motor neurons of *Smn^2B/-^* mice [57]. However, given that p53 induction alone is necessary but not sufficient to drive death of motor neurons *in vivo* [27,28], we looked for amino-terminal phosphorylation of p53 that we showed to be a prerequisite for activation of the neurodegenerative process [27,29]. Specifically, we investigated the phosphorylation of p53 at serine 18, which we have previously shown to be a p38αMAPK-mediated event that marks vulnerable SMA motor neurons destined to die in *SMNΔ7* mice [29]. Importantly, we found no evidence for p53^S18^ phosphorylation in lumbar SMA motor neurons from *Smn^2B/-^* mice on FVB/N background. The absence of this post-translational modification of p53 may explain why motor neuron survival is unaffected in this mouse model and corroborates our proposed model in which convergence of upregulation and amino-terminal phosphorylation of p53 are distinct events necessary for driving motor neuron death [27–29]. Despite contrasting observations [55,56], which may be related to issues of reliability in motor neuron counting described above, this conclusion is also consistent with recent findings that the modest loss of L1 motor neurons in *Smn^2B/-^* mice on C57BL/6 background correlates with the expression of phospho-p53^S18^ and is rescued by p53 inhibition [37].

In sum, our work highlights the importance of monitoring the same pathogenic events with the same experimental approaches when comparing sensory-motor circuit pathology in different mouse models of SMA. Together with previous studies [30,36,37,45–47], we identify the loss of proprioceptive synapses on motor neurons as a conserved cellular defect induced by SMN deficiency across mouse models of SMA. Similarly, NMJ denervation of axial muscles such as the QL displays good conservation across models and accurately reflects the proximo-distal gradient of muscle vulnerability characteristic of SMA patients. In contrast, selective loss of motor neurons unexpectedly emerges as the most distinguishing feature across mouse models of SMA despite being a hallmark of the human disease. While the *SMNΔ7* model shows early onset and progressive death of specific motor neuron pools [27,30], in *Smn^2B/-^* mice the same motor neurons are either entirely spared (FVB/N background, this study) or a subset thereof only affected at very late disease stages (C57BL/6 background, [37]). Moreover, no loss of motor neurons has recently been reported in the *Taiwanese* SMA model [37]. Thus, not all mouse models of SMA are equally poised for the study of every aspect of sensory-motor circuit pathology. In this context, *SMNΔ7* mice rather than *Smn^2B/-^* and *Taiwanese* mice are better suited for *in vivo* testing of neuroprotective drugs that selectively target the motor neuron death pathway. Collectively, these findings should help guide the selection of the most appropriate mouse models for elucidating specific disease mechanisms and pre-clinical testing of SMN-independent therapies.

## Acknowledgements

We are grateful to George Mentis and Christian Simon for comments and critical reading of the manuscript. We thank Rashmi Kothary for providing the *Smn^2B^* mouse line. This work was supported by NIH grant R01NS116400 (L.P.) and the NSF Graduate Research Fellowship Program (M.T.). The funders had no role in study design, data collection and analysis, decision to publish, or preparation of the manuscript.

## Author Contributions

L.P. conceived and supervised the study. M.J.C. and M.K.T. performed the experiments. M.J.C., M.K.T. and L.P. analyzed the data. M.J.C. and L.P. wrote the manuscript.

---

**Fig S1.**
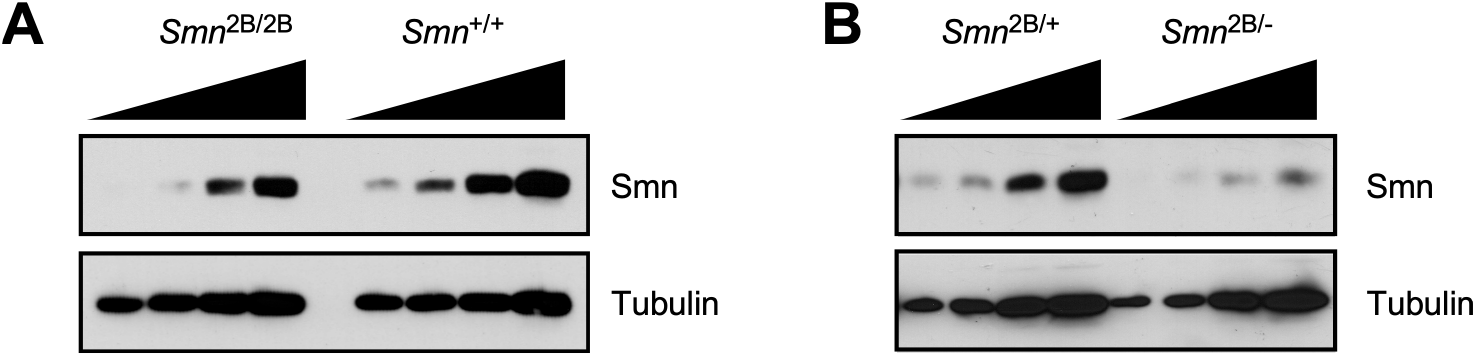
Analysis of Smn expression from the *Smn^2B^* allele in the mouse spinal cord. (A) Western blot analysis of Smn levels in the spinal cord from *Smn^+/+^* (wild type) and *Smn^2B/2B^* mice at P16. (B) Western blot analysis of Smn levels in the spinal cord from *Smn^2B/+^* an *Smn^2B/-^* mice at P16. Two-fold serial dilutions of equal amounts of extracts are shown. Tubulin was probed as a loading control.

**Fig S2.**
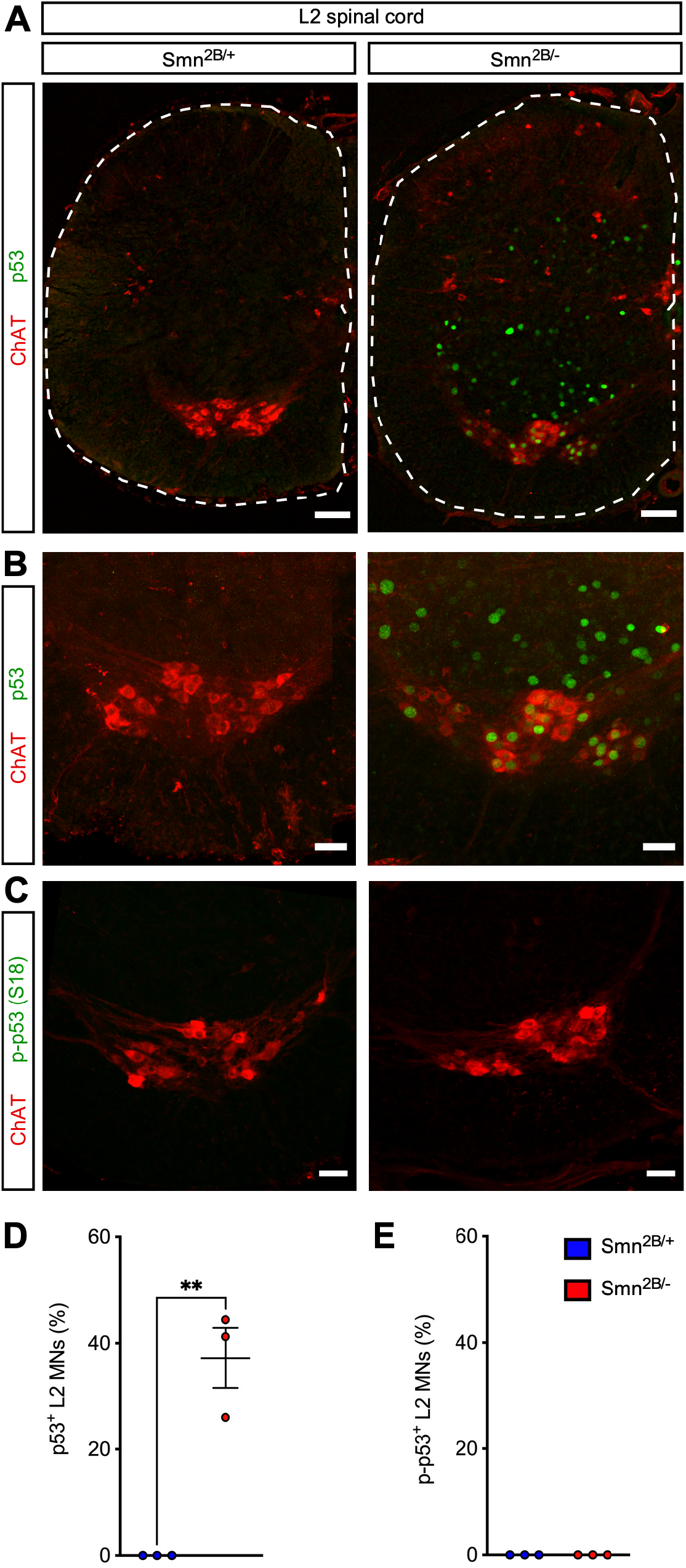
Smn deficiency induces p53 accumulation but not serine 18 phosphorylation in L2 motor neurons of *Smn^2B/-^* SMA mice. (A) ChAT and p53 immunostaining of the L2 spinal cord from control (*Smn^2B/+^*) and SMA (*Smn^2B/-^*) mice at P16. Scale bars: 100 μm. (B) ChAT and p53 immunostaining of L2 motor neurons from the same groups as in (A). Scale bars: 50 μm. (C) ChAT and phospho-p53^S18^ immunostaining of L2 motor neurons from the same groups as in (A). Scale bars: 50 μm. (D) Percentage of p53^+^ L2 motor neurons from the same groups as in (A). (E) Percentage of phospho-p53^S18+^ L2 motor neurons from the same groups as in (A). Data represents individual values, mean and SEM from 3 mice per group. Statistics were performed with two-tailed unpaired Student’s t-test. ** P < 0.01.

**Fig S3.**
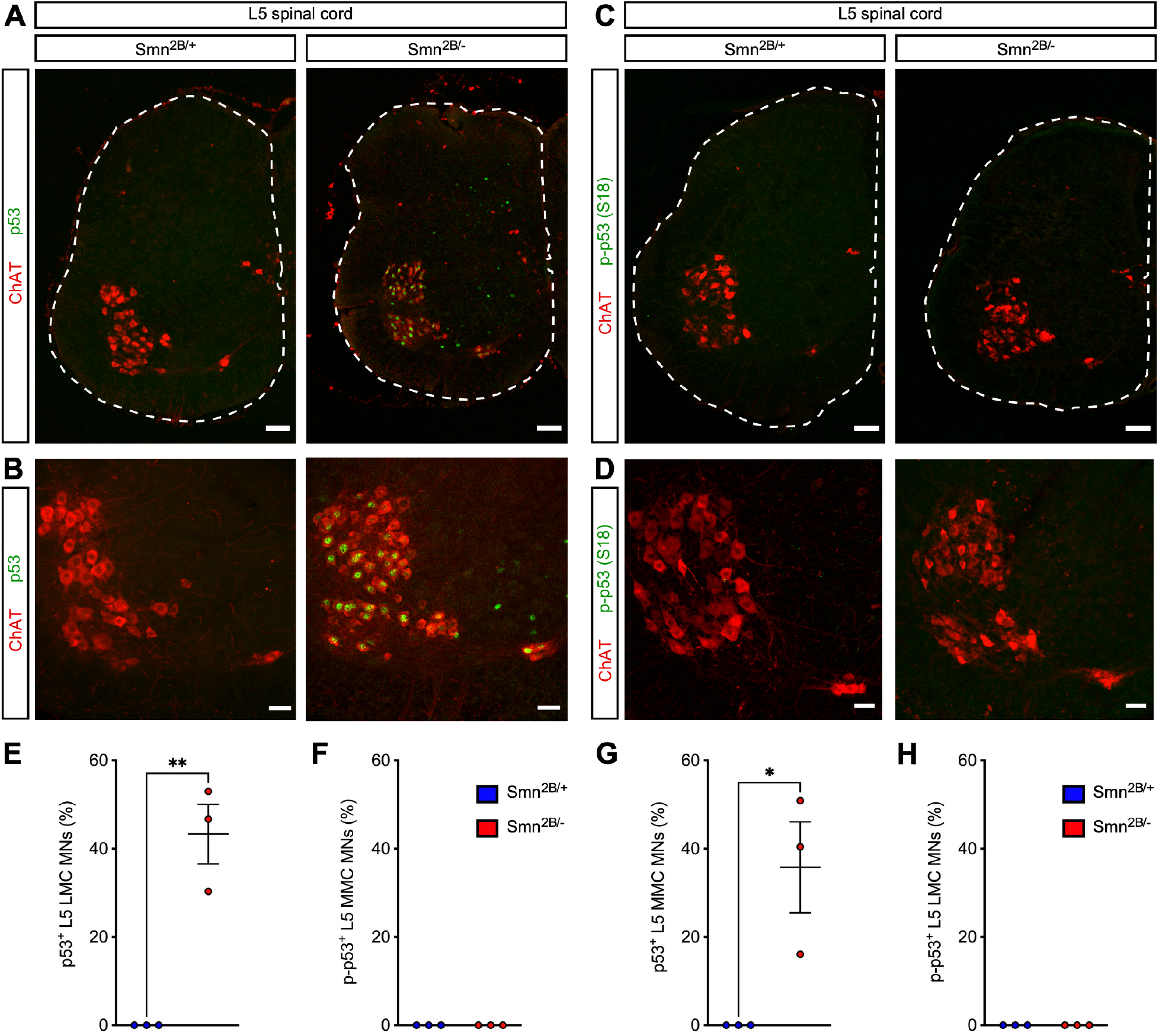
Smn deficiency induces p53 accumulation but not serine 18 phosphorylation in L5 motor neurons of *Smn^2B/-^* SMA mice. (A) ChAT and p53 immunostaining of the L5 spinal cord from control (*Smn^2B/+^*) and SMA (*Smn^2B/-^*) mice at P16. Scale bars: 100 μm. (B) ChAT and p53 immunostaining of L5 motor neurons from the same groups as in (A). Scale bars: 50 μm. (C) ChAT and phospho-p53^S18^ immunostaining of the L5 spinal cord from the same groups as in (A). Scale bars: 100 μm. (D) ChAT and phospho-p53^S18^ immunostaining of L5 motor neurons from the same groups as in (A). Scale bars: 50 μm. (E and G) Percentage of p53^+^ L5 LMC (E) and L5 MMC (G) motor neurons from the same groups as in (A). (F and H) Percentage of phospho-p53^S18+^ L5 LMC (F) and L5 MMC (H) motor neurons from the same groups as in (A). Data represents individual values, mean and SEM from 3 mice per group. Statistics were performed with two-tailed unpaired Student’s t-test. ** P < 0.01; * P < 0.05.

